# Association between Brain Structure and Cognitive Ability during Adolescence: Insights from a Comprehensive Large-Scale Analysis of 9 to 15 Year-Olds

**DOI:** 10.1101/2024.06.18.599653

**Authors:** Jiadong Yan, Yasser Iturria-Medina, Gleb Bezgin, Paule Joanne Toussaint, Ke Xie, Liang He, Judy Chen, Kirsten Hilger, Erhan Genç, Alan Evans, Sherif Karama

## Abstract

Significant changes occur in brain structure and cognitive abilities during adolescence. Investigating their association can provide insight into brain-based cognitive development, yet previous studies were limited by narrow brain measures, small samples, and lacking focus on age-related variation. Here, we analyzed a large cohort (*N* = 8,534, age 9–15) from the Adolescent Brain Cognitive Development dataset. Using structural MRI and diffusion imaging, we derived 16 regional structural measures and integrated them via morphometric similarity networks to characterize 16,563 regional, connectivity, and hub features. We applied large-scale computational models to investigate their associations with performance on seven cognitive subtests and general intelligence (*g*), as well as age-related changes. Brain areas most strongly associated with cognitive ability also showed the greatest age-related variability in these associations, located primarily in the frontal, temporal, and occipital lobes. Structural MRI measures exhibited stronger associations with cognition and greater age-related variability than diffusion-derived metrics, while global hub measures showed stronger and more variable associations than local measures. Overall, our study provides a comprehensive and reliable understanding of brain structure-cognition associations during adolescence.

## Introduction

Mounting evidence highlights the intricate relationship between brain structure and cognitive ability^1–6^. Since both brain structure and cognitive ability change significantly during adolescence^7–9^, a comprehensive study of their association is crucial for understanding brain development and its impact on cognitive maturation. Numerous studies have shown that region-wise brain structural measures are significantly correlated with performance on various cognitive tests^6,10^. For instance, structural Magnetic Resonance Imaging (sMRI) measures like regional cortical thickness, surface area, and gray matter volume are significantly correlated with cognitive performance^7,11–14^. Moreover, variations in regional cortical gyrification^15,16^ and region-wise gray matter microstructure from Diffusion-Weighted Imaging (DWI)^17–19^ have also been linked to differences in cognitive ability.

Understanding brain structure-cognition associations requires examining not only regional structure but also the interconnections between regions^3,20,21^. Previous studies have used DWI to identify patterns of fiber connections significantly associated with cognitive ability^1,19,22^. Moreover, structural connections characterized by using morphometric similarity networks (MSNs) have also been linked to cognitive test performance^21,23^. For instance, a recent study reported that weaker MSN-based cortico-subcortical connections are associated with higher cognitive ability^23^. Highly central brain areas, referred to as hubs^24^, play a crucial role in cognitive ability^4,25–28^. Consistent with this, the Parieto-Frontal Integration Theory (P-FIT) of intelligence emphasizes that communication between specific brain hubs is pivotal for cognitive functioning^29,30^. The organization of structural brain networks, including properties like modular segregation derived from DWI, is also linked to cognitive test performance^31–33^, a finding supported by studies of structural covariance networks (SCNs)^34^. Finally, the nodal degree of MSN hubs, a measure of a region’s overall connectedness, is highly correlated with both verbal and non-verbal cognitive performance^21^.

Although previous research has explored the association between various brain structural measures and cognitive abilities, several key limitations persist. No single study has simultaneously combined these diverse measures into one computational model to create a comprehensive representation of how brain architecture relates to cognitive differences. Moreover, the use of different data samples and analysis methods across studies has produced heterogeneous effects with limited comparability. Sample size remains another critical limitation; many studies include fewer than 500 participants. While some recent studies using the Adolescent Brain Cognitive Development (ABCD) dataset have incorporated thousands of participants^14,23,35^, they typically use narrow age ranges (e.g., 9–11 years) that fail to capture most of adolescent period, which can introduce sampling error and lead to an overestimation of effects. Finally, the temporal stability of these associations is an important open question. Although both brain structure and cognitive ability undergo dramatic changes during this developmental period, the effect of age on brain structure-cognition associations has not been extensively investigated^36–38^.

To address these research gaps, we obtained a comprehensive set of brain structural measures from the ABCD dataset. The dataset’s multimodal brain images, including sMRI and DWI^39^, enabled us to calculate comprehensive region-wise cortical and subcortical brain structural measures^40^. From these regional measures, we used morphometric similarity networks (MSNs) to characterize the structural connections between each pair of brain regions^21^ and then calculated graph theoretical measures to capture the hub-wise characteristics of connected brain regions^41^.

MSNs were used for their ability to characterize structural connections and networks by calculating measurement similarities across brain regions, which can reflect similarities in cytoarchitecture, gene expression, and white matter connections^21,42^. Variations in MSN topologies have been shown to correlate with age, cognitive function^43^, and mental health disorders^44^. Compared to DWI, which may suffer from under-recovered long-distance projections and false-positive connections, and SCNs, which rely solely on a single measure, MSNs are suggested to be more comprehensive and accurate^21^. These approaches enabled us to establish a comprehensive whole-brain structure set, including region-wise, connection-wise, and hub-wise measures. To assess associations between these measures and cognitive ability, we developed a robust large-scale computational model based on previous work^45^. This framework allows the structural inputs to control for one another, yielding comparable results across different inputs. To address limitations related to small sample sizes, we included 8,534 participants from the ABCD project^39^. Finally, we extended this framework to investigate how brain structure-cognition associations change with age during adolescence. The overall flowchart of the association analyses was shown in **Fig. 1**.

**Fig. 1.**
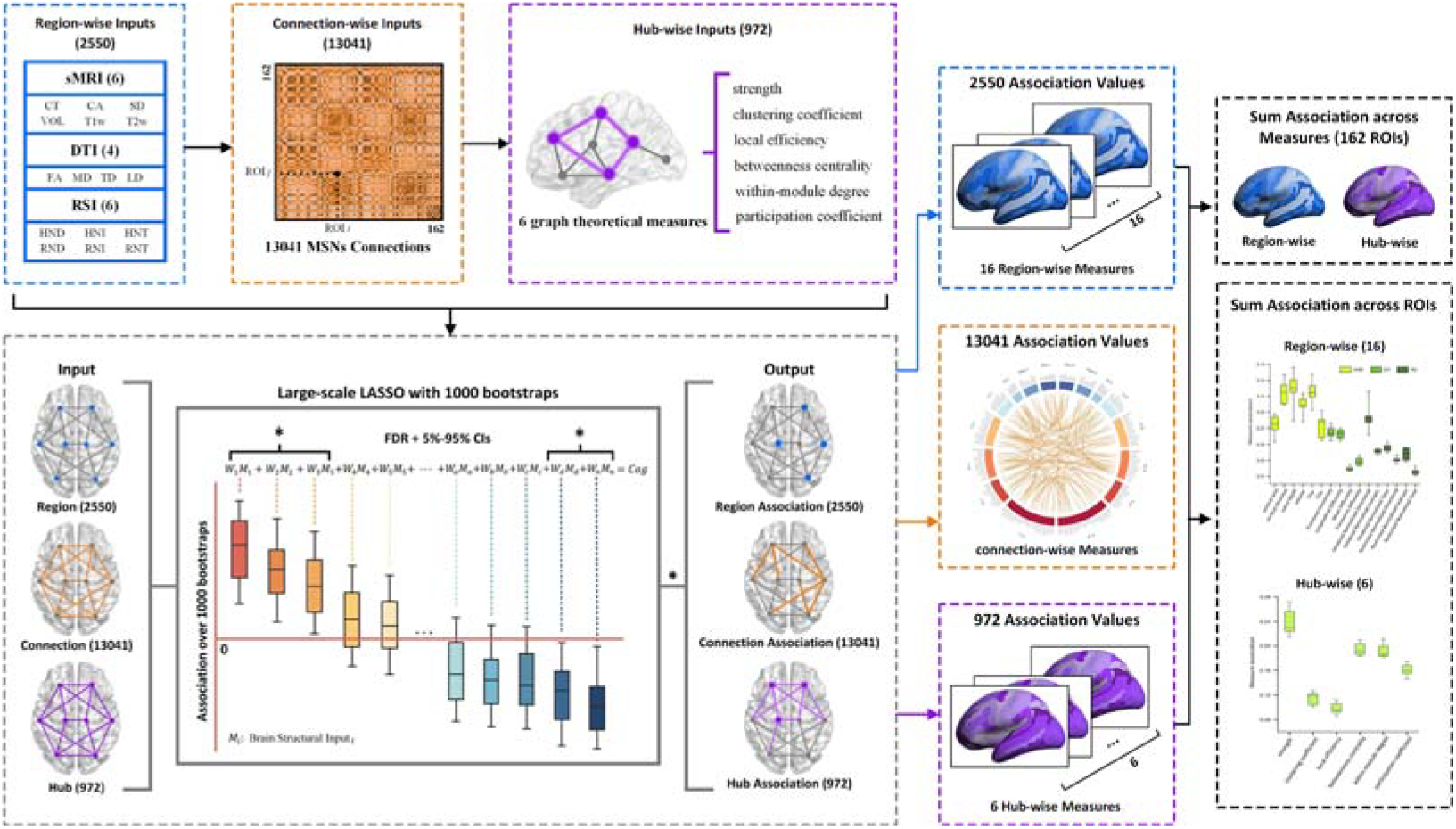
Flowchart of the association analysis. The analysis framework was used to investigate both specific and summary associations.

This study had three primary objectives. First, we sought to identify the brain regions, connections, and hubs with the strongest associations with cognitive ability. Second, we investigated which brain structural features—among sMRI, Diffusion Tensor Imaging (DTI), Restriction Spectrum Imaging (RSI), and graph theoretical hub measures—were most associated with cognition. Third, we examined the temporal variability in these brain structure-cognition associations. Overall, this work aimed to provide a comprehensive and reliable understanding of the relationship between brain structure and cognitive ability during adolescence.

## Results

Using a large ABCD sample and a comprehensive collection of brain structural measures, we employed robust computational models to investigate associations between brain structure and cognitive ability, as well as their temporal stability during adolescence. First, we identified the brain regions, connections, and hubs with the strongest associations with cognitive ability. Second, to identify the whole-brain summary measures most associated with cognition, we summed the region-and hub-wise associations across all 162 ROIs. A similar summary analysis was not applicable to structural connections, as each was represented by a single value. Third, we investigated the age-related stability of these brain-cognition associations across all structural properties. Finally, we assessed the effectiveness and robustness of our model using two validation methods.

### Associations between *g* and brain regions, connections, and hubs

After identifying significant brain structural inputs associated with cognitive ability, we investigated the summary association values for brain regions and hubs, as well as the individual association values for brain connections. **Figure 2** shows associations with general intelligence (*g*), while associations with the other seven cognitive subtests are presented in **Supplementary Table 2**. For region-and hub-wise results, we calculated the sum of absolute association values across all structural measures (sMRI, DTI, RSI, and graph theoretical hub features) for each ROI and represented these on brain maps. To ensure comparability between cortical regions (16 measures) and subcortical regions (13 measures), we adjusted the summary values for subcortical ROIs by multiplying them by 16/13. For connection-wise results, the top 1% of connections are displayed in a connectogram in **Figure 2** (results for other top *p*% thresholds are in **Supplementary Fig. 1**). Furthermore, for between-lobe comparisons that account for variations in the number of ROIs per lobe, we counted the number of regions, connections, and hubs within the top *p*% of association values. We examined thresholds from 1% to 30% and found 20% to be representative (**Supplementary Materials S1, Supplementary Fig. 2**); these results are shown in **Fig. 2B**, with others presented in **Supplementary Fig. 3**. Overall, the results highlighted the frontal, temporal, and occipital lobes as having the strongest associations with cognitive ability across regional, connection, and hub measures. Conversely, the insular cortex demonstrated a relatively low association. While subcortical nuclei were weakly associated in the connection-and hub-wise analyses, they were highly associated in the region-wise results, especially the right putamen and caudate.

**Fig. 2.**
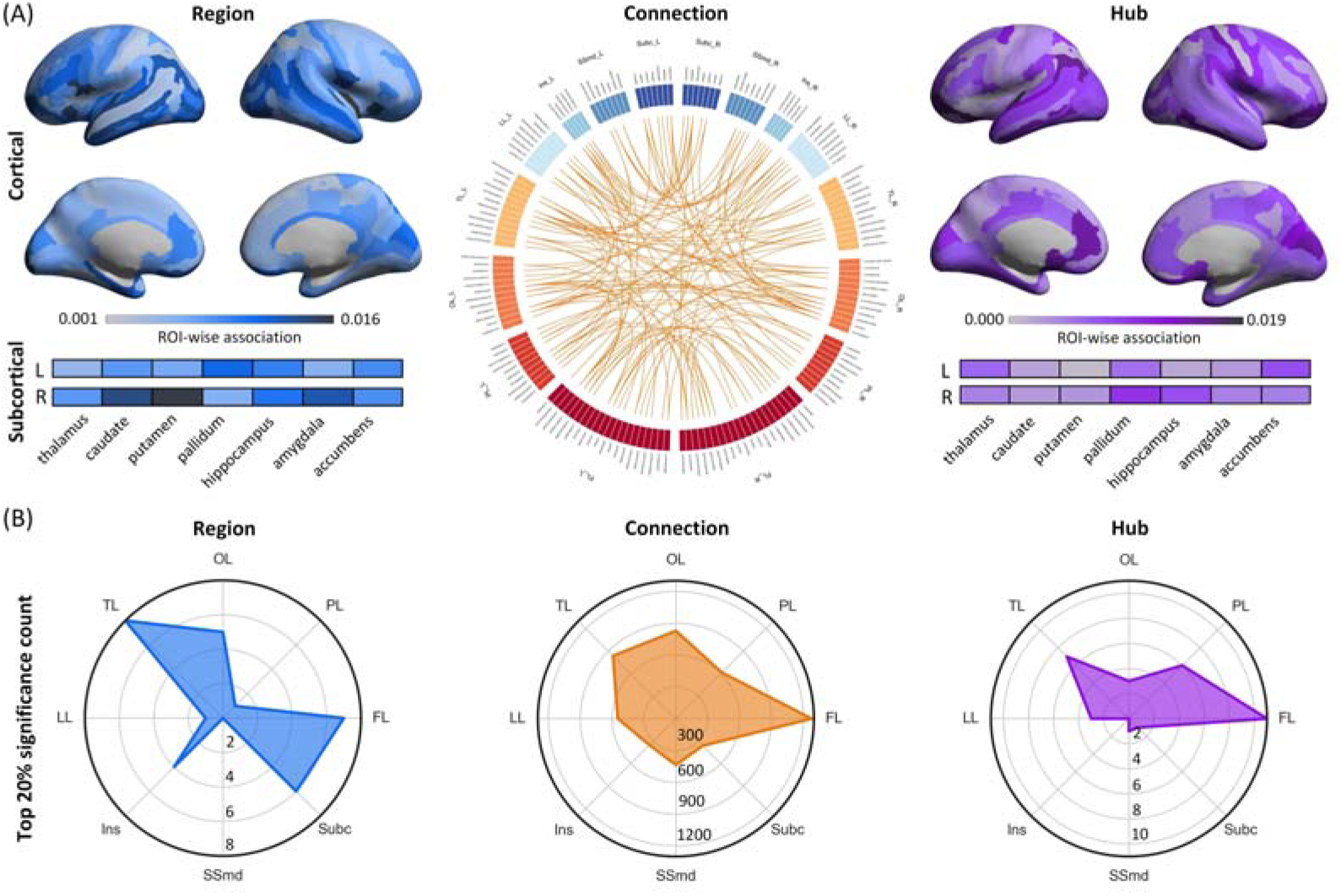
Associations between brain structure and adolescent *g* across different brain regions, connections, and hubs. Blue, orange, and purple represent region-wise, connection-wise, and hub-wise results, respectively. (A) Associations for brain regions, connections (top 1% shown), and hubs. For a list of the 162 ROIs and their lobe categories, see **Supplementary Table 1**. A high-resolution version of the connectogram is available in **Supplementary Fig. 1**. (B) The number of brain regions, connections, and hubs within the top 20% of association values is shown for each lobe.

**Fig. 3.**
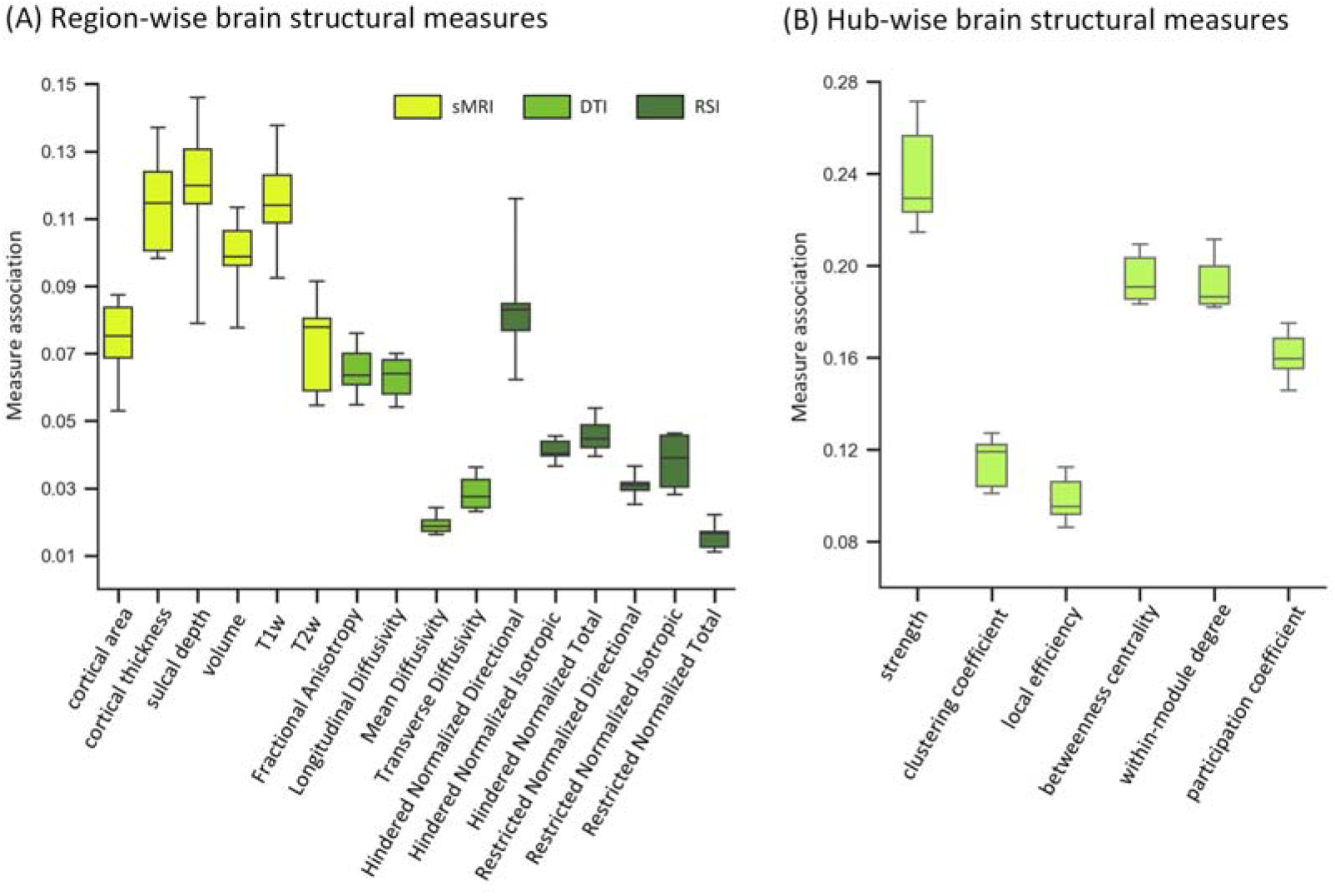
Associations between whole-brain summary measures and adolescent cognitive ability. (A) Associations for summary region-wise measures. (B) Associations for summary hub-wise measures.

### Associations between whole-brain summary measures and cognitive ability

For each of the 22 summary whole-brain measures (16 region-wise and 6 hub-wise), we calculated the sum of absolute association values across all ROIs to represent the measure’s overall association with cognitive ability. Because three sMRI measures (cortical surface area, thickness, and sulcal depth) were calculated for cortical ROIs only, we adjusted their corresponding summary values by multiplying by 162/148 to ensure a fair comparison. The region-wise results (**Fig. 3A**) show that sMRI measures had stronger associations with cognitive ability than DTI and RSI measures, although FA, LD, and HND also showed relatively high associations. For the summary hub-wise measures (**Fig. 3B**), local network attributes (clustering coefficient and local efficiency) were less associated with cognitive ability than global attributes (strength, betweenness centrality, within-module degree, and participation coefficient), with strength showing the strongest association.

### Age-related instability in associations between *g* and brain regions, connections, and hubs

To quantify the temporal instability of brain-cognition associations, we used the absolute value of the age interaction term from **Equation 2** in the Methods section. For region-and hub-wise results, we summed these absolute interaction values across all structural measures for each ROI, again adjusting the values for subcortical ROIs (multiplying by 16/13) to ensure comparability. For connection-wise results, the absolute interaction value itself represented the instability. Results for the other seven cognitive subtests are available in **Supplementary Table 2**. As in the previous section, **Fig. 4A** shows the connection-wise results (top 1% shown), and **Fig. 4B** shows the lobe distribution for the top 20% of unstable regions, connections, and hubs. Results for other top *p*% thresholds are in **Supplementary Fig. 4** and **Supplementary Fig. 5**. These analyses highlighted the occipital, frontal, and temporal lobes as exhibiting the greatest age-related change in their brain-cognition associations. Conversely, the insular cortex demonstrated the highest stability. Combining these findings with the results in **Fig. 2**, we found that lobes with stronger associations with cognitive ability also tended to have more unstable associations during adolescence.

**Fig. 4.**
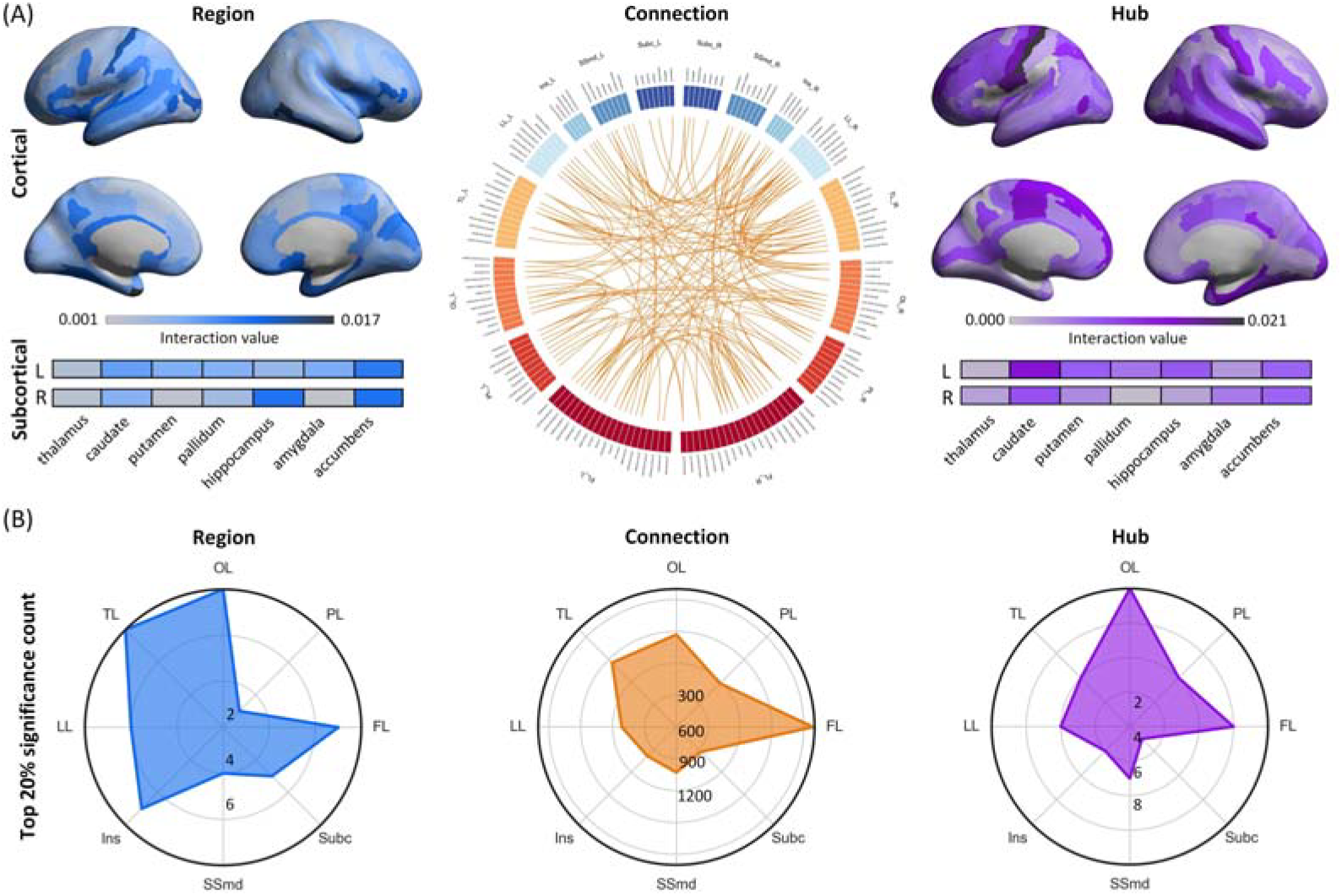
Age-related instability in the associations between brain structure and adolescent *g*. Blue, orange, and purple represent region-wise, connection-wise, and hub-wise results, respectively. (A) Instability in the associations for brain regions, connections (top 1% shown), and hubs. A high-resolution version of the connectogram is available in **Supplementary Fig. 1**. (B) The number of the top 20% most unstable brain regions, connections, and hubs in their brain-cognition associations during adolescence, shown within each lobe.

**Fig. 5.**
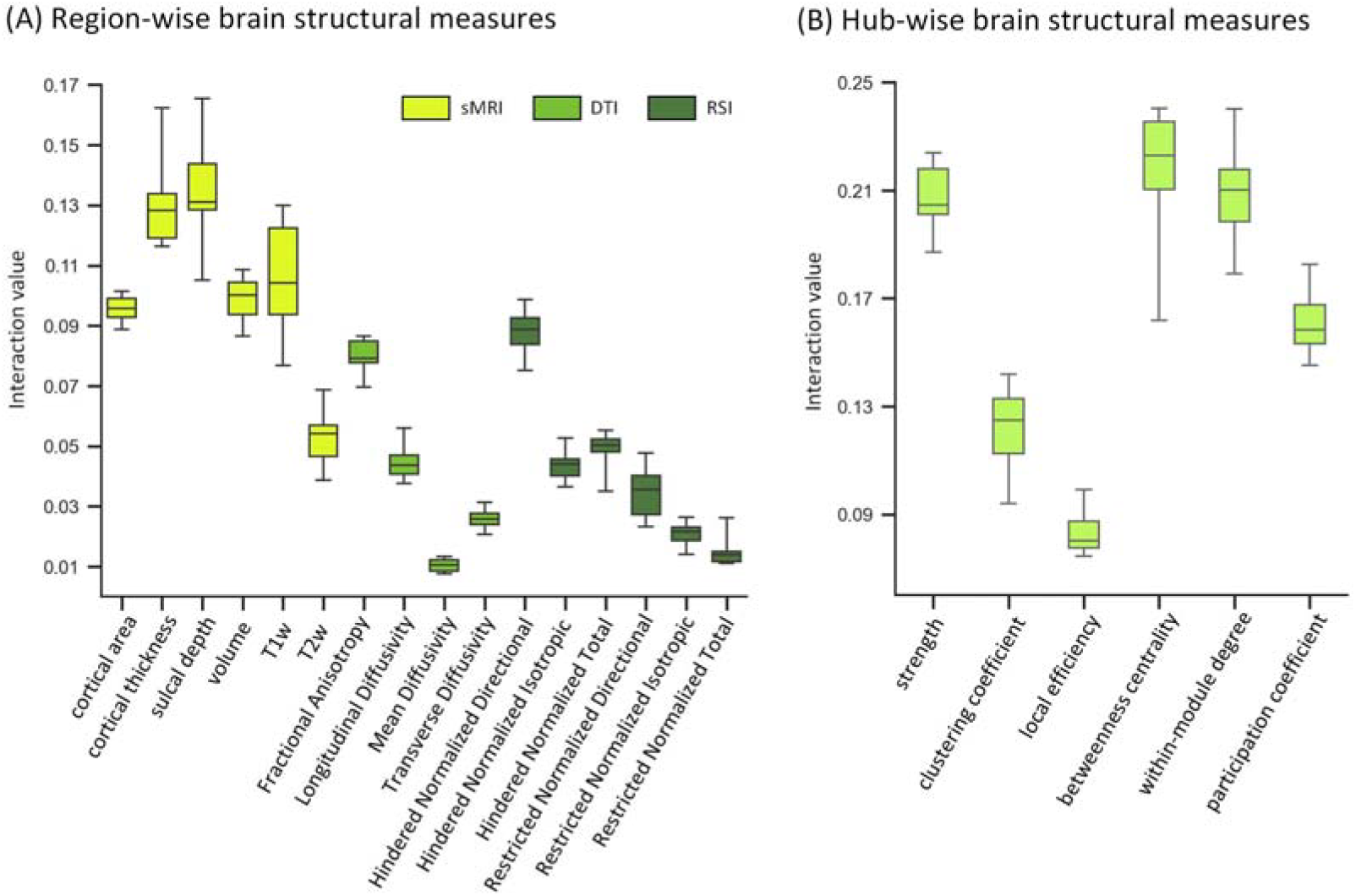
Age-related instability in the associations between whole-brain summary measures and cognitive ability. Higher interaction values indicate greater instability in the corresponding associations. (A) Instability for whole-brain summary region-wise measures. (B) Instability for whole-brain summary hub-wise measures.

### Age-related instability in associations between whole-brain summary measures and cognitive ability

For each of the 22 whole-brain summary measures, we calculated the sum of the absolute interaction values across all ROIs to represent the age-related instability of its association with cognitive ability. As in the previous section, we adjusted the summary values for the three cortical-only sMRI measures (CA, CT, and SD) by multiplying by 162/148 to ensure a fair comparison. The region-wise results (**Fig. 5A**) showed that associations involving sMRI measures were more unstable with age than those involving DTI or RSI measures, although FA in DTI and HND in RSI also displayed relatively high age-related instability. For the summary hub-wise measures (**Fig. 5B**), local network attributes (clustering coefficient and local efficiency) were more stable than global attributes (strength, betweenness centrality, within-module degree, and participation coefficient), with local efficiency being the most stable. Combining these observations with the results in **Fig. 3**, we found that summary brain measures with stronger associations with cognitive ability also tended to have more unstable associations during adolescence.

### Effectiveness and robustness of the utilized large-scale computational model

To evaluate the large-scale computational model used for investigating brain structure-cognition associations, we assessed its effectiveness and robustness. To assess effectiveness, we calculated the coefficient of determination (*R*^*2*^) between the observed and estimated cognitive scores for 1,000 bootstraps. We then averaged these 1,000 *R*^*2*^ values to quantify how well the model captured the brain-structure associations for each cognitive score. For each of the eight cognitive measures (seven subtests and *g*), we obtained six mean *R*^*2*^ values from models (1) and (2) across the region, connection, and hub inputs (*i*.*e*., 2 × 3 = 6; **Fig. 6A**). Mean *R*^*2*^ values ranged from 0.12 to 0.63, depending on the cognitive ability examined. The strongest mean *R*^*2*^ was observed for the association between brain structure and *g*. To assess robustness, we varied the number of bootstraps used in the model. For all 48 models (3 input types × 8 cognitive measures × 2 model types), we conducted iterations with 50, 100, 200, 300, 400, and 500 bootstraps, repeating the process 100 times for each number. We then calculated the average correlation among the 100 repetitions for each bootstrap number as a measure of robustness (**Fig. 6B**). The results showed that with 400 bootstraps, the correlations consistently exceeded 0.979. With 500 bootstraps, all correlation values exceeded 0.983, averaging 0.993. These high values indicate strong model reliability, validating our use of 1,000 bootstraps in the main analyses.

**Fig. 6.**
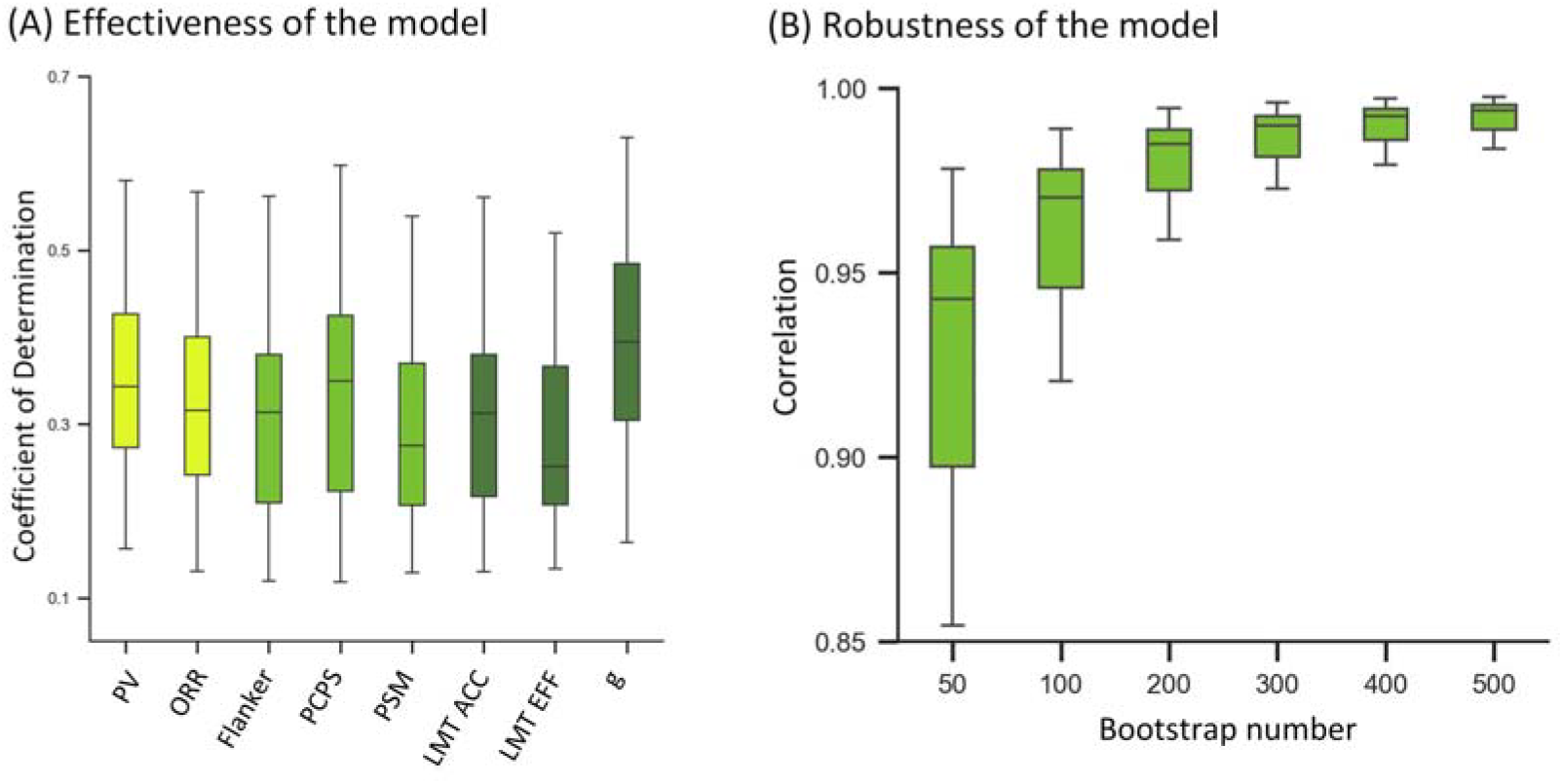
Validation of results. (A) The effectiveness of the model. (B) The robustness of the model.

## Discussion

Using data from 8,543 participants aged 9–15 years from the ABCD 5.1 release, we conducted a comprehensive analysis of the association between brain structure and cognitive ability during adolescence. We found that the brain areas most strongly associated with cognitive ability were located primarily in the frontal, temporal, and occipital lobes, whereas subcortical regions and the insular cortices showed the weakest associations. Notably, the brain areas most strongly associated with cognition also demonstrated the largest age-related changes in these associations. Among the different structural features examined, sMRI measures had the strongest associations with cognitive ability and the greatest age-related changes compared to DTI and RSI measures. Finally, hub-wise measures of global network attributes were more strongly associated with cognitive ability and showed greater age-related change than measures of local network attributes. Importantly, our large-scale computational model demonstrated promising effectiveness and robustness, suggesting its potential for future brain-behavior research.

### Cognitive associations with brain regions, connections, and hubs

Our region-wise analysis showed that the frontal lobe, temporal lobe, and subcortical regions (especially the right putamen and caudate) contained the brain regions most strongly associated with cognition, a finding consistent with previous work. For instance, adolescent development of regional GABAergic interneuron subtypes in the prefrontal cortex (PFC) contributes to the improvement of higher cognitive functions like cognitive flexibility^37^. Speculatively, these GABAergic interneurons might be reflected in our brain structural measures, thereby contributing to the association we observed between the frontal lobe and general intelligence (*g*). Similarly, regions within the temporal lobe, such as the anterior superior temporal sulcus and superior temporal gyrus, are significantly activated by spoken words and abstract concepts^46^. This suggests that the regional structural differences we observed in the temporal lobe might underlie individual differences in language-related cognitive abilities. Regarding the putamen and caudate nuclei, previous studies have shown that structural lesions in these areas impair delayed response, alternation behavior, and visual discrimination learning^47,48^. Our results therefore support the crucial role of the putamen and caudate nuclei in cognitive ability.

Our connection-wise analysis revealed that the structural connections of the frontal, occipital, temporal, and parietal lobes were most strongly associated with cognition. These lobes encompass nearly the entire cerebrum, corroborating recent proposals that effective brain-wide connections form the foundation of cognitive function^27,49^. Additionally, this finding aligns with the Parieto-Frontal Integration Theory (P-FIT), suggesting that enhanced communication between multiple brain lobes, especially the parietal and frontal lobes, leads to improved cognitive ability during adolescence^29^. Our hub-wise analyses showed that most cognition-associated hubs were located in the frontal lobes. This is compatible with previous work suggesting that the PFC functions as a central hub that mediates cognitive control, while its cytoarchitectonic heterogeneity helps coordinate and integrate different tasks^50^. Given that MSNs reflect similarities in cell-type diversity, a feature associated with stronger structural connections^21^, this cellular information may be captured by our MSN-based hub characteristics, thus contributing to the observed association with cognitive abilities. Brain areas in the parietal lobe also showed relatively strong associations, which might be partly due to their integrative role in facilitating communication across the brain network during adolescence^51,52^.

Regarding age-related changes in the brain-cognition association during adolescence, brain areas in the occipital lobe were the most unstable. While an early study showed that the cortical gray matter volume of the occipital lobe continued to increase up to 20 years of age^9^, more recent work with strict quality control of structural MRI data has revealed that both cortical thickness and surface area in the occipital lobe undergo substantial developmental changes throughout adolescence^8,53^. Moreover, the occipital lobe is significantly activated when performing many cognitive tasks, especially those related to visuospatial processing^54^ and is one of the most highlighted brain structures in the P-FIT model^29,30^. These two observations may explain the instability in the association between occipital structure, such as gray matter volume, and cognitive abilities, such as visuospatial ability. In addition, regions in the frontal lobe showed relatively high age-related changes, which might be due to the rapid development of its local information processing capabilities and enhanced communication with regions in other lobes to support higher-order cognitive abilities during adolescence^55,56^. Interestingly, brain areas showing stronger associations with cognitive abilities also tended to show the greatest change in the strength of their associations with age. This finding aligns with previous studies that found significant changes during childhood and adolescence in both the brain structure of the most associated lobes (i.e., frontal, temporal, and occipital) and their corresponding cognitive abilities^57–59^.

### Cognitive associations with whole-brain summary measures

Among the 16 summary region-wise measures we examined, cortical thickness, sulcal depth, and T1w density emerged as the three measures most strongly associated with cognitive ability. This is in line with previous research revealing significant associations between these three measures and cognition. Most variability in cortical thickness is accounted for by the amount of glial and capillary support, neuronal density, and the level of dendritic arborization, which has been suggested to provide the cellular foundation for human cognitive ability^6,7,12,13^. Regarding sulcal depth, previous studies found that only the gyrification of six-layered cortical regions was significantly related to intelligence^15,16^. Although primary sensory and motor cortices also exhibit cortical folding, they differ in cellular structure, especially in having a reduced or absent layer IV. These regions were not among those associated with intelligence. This pattern suggests that differences in laminar organization and cell-type composition may contribute to the observed association between gyrification and cognitive ability. Additionally, the significance of T1w density may stem from its indirect reflection of the structural aspects of neurons, glial cells, and synaptic connections in brain gray matter, which are believed to be crucial for cognitive processes^60–63^.

Importantly, we observed that the significance of sMRI measures is notably higher than that of DTI and RSI measures. A possible reason is that gray matter morphometry from sMRI might be more reflective of cellular information than the microstructural features captured by DTI and RSI measures^15,64^. Alternatively, many DTI and RSI measures may be contaminated with noise during data acquisition and metric calculation compared to sMRI, thereby weakening the association^65,66^. Moreover, DTI and RSI might not be optimal for gray matter, although models like neurite orientation dispersion and density imaging (NODDI) perform well in this tissue^17^. Nonetheless, two DTI measures (FA and LD) and one RSI measure (HND) also showed relatively high associations, comparable to sMRI measures. Higher FA values typically indicate more organized and intact neural fiber structures, while reduced LD may suggest damage or degeneration within neural fiber bundles^26^. HND primarily reflects the restricted diffusion of water molecules along neural fiber bundles, providing insights into fiber orientation and microstructural integrity^67,68^. Although DTI and RSI measures were generally less associated with cognition, specific DWI measures (FA, LD, and HND) characterizing fiber directionality and microstructural integrity still showed a high association. This might indicate that these microstructural characteristics significantly affect cognitive ability. Finally, we found that summary hub-wise measures related to local graph theoretical attributes were less associated with cognition than those related to global attributes. This finding again supports the notion that cognitive ability arises more from brain-wide communication rather than from the connectedness of local brain regions^4,25,27^.

Regarding the age-related changes in brain-cognition associations, summary brain measures that were more associated with cognition also tended to be more unstable. Many previous studies have reported significant changes in brain gray matter during adolescence, which can be reflected in sMRI measures. For instance, synaptic density within gray matter increases during early life across the whole brain, then decreases in the somatosensory cortex during childhood and in the prefrontal cortex during adolescence^69,70^. This structural developmental trajectory may be linked to the instability of related sMRI measures, such as CT and T1w density, in their association with cognition during this period. In terms of microstructural features, increasing FA in the inferior longitudinal fasciculus and inferior fronto-occipital fasciculus has been observed in adolescence^70,71^. Similarly, key properties of structural brain networks, such as nodal degree (related to strength) and modularity (related to within-module degree and participation coefficient), also undergo significant reorganization during this period^72^. Together, these studies indicate that the brain summary measures most associated with cognition all exhibit rapid and regionally heterogeneous developmental changes, which may explain the high temporal variability and unstable patterns of brain-cognition associations during adolescence.

### Cognitive abilities vary in their brain structural associations

Our results showed that all 7 cognitive performance measures and the latently modeled *g* were significantly associated with the brain structural measures employed here. Among these cognitive measures, *g* demonstrated the highest association, while cognitive performance scores related to episodic memory, attention, visuospatial ability, and reading expressed relatively lower associations. As *g* captures the shared variance among cognitive subtest scores^73,74^, it is plausible to assume that *g* links to more common and stable brain structural measures. In terms of episodic memory, several studies revealed its rapid development during childhood and adolescence^75,76^. Specifically, these studies suggest that during early childhood, episodic memory relies on hippocampal binding mechanisms, while further improvements result from the development of the prefrontal cortex. This transition might cause instability in the associated brain measures from ages 9 to 15, which would explain the lower brain-cognition association value observed in our data-driven analysis. With respect to attention, previous studies found that its development followed sexually divergent trajectories during adolescence^77,78^. This may indicate a complex mechanism of sex differences in the development of Flanker attention ability, potentially causing a non-linear relationship between brain structure and attention. In terms of visuospatial and reading abilities, previous studies found highly significant effects of age on their development during adolescence^79,80^, which might also lead to a lower brain-cognition association value. Overall, previous studies suggest a high complexity and variability of brain-cognition associations. However, when investigating the association between brain structure and *g*, the shared brain features of different cognitive tasks might be more stable across these complex mechanisms, thus contributing to the higher association observed for *g*.

### Potential of our model for future research on brain-behavior associations

A key advantage of this study is its use of machine learning methods on a large sample with a comprehensive set of brain structural measures. This combination enabled us to obtain detailed and robust results on brain structure-cognition associations and to compare the association values for each measure and brain area while controlling for all others within a single model. However, using a large number of input features relative to the number of subjects (the *N/P ratio* problem) can lead to unreliable interpretations in traditional machine learning methods^81^. We therefore chose the LASSO method to mitigate the *N/P ratio* problem^82^. While effective, LASSO’s greedy approach to parameter optimization can lead to biased interpretations when inputs are multicollinear, as is the case with brain structural measures. To address this limitation, we introduced a feature random optimization strategy, which randomizes the order of parameter computation across 1,000 bootstraps. By averaging the results, this approach counteracts the weaknesses of LASSO and reduces bias in model interpretations, a strength confirmed by our model’s robustness test. The observation that reliable results were achievable with as few as 200 bootstraps suggests that our approach can be readily transferred to future studies investigating multivariate brain-behavior associations, even when input variables have high collinearity or face the *N/P ratio* problem.

Although our in-sample R^2^ for brain–cognition associations was relatively high, competitors in the ABCD Neurocognitive Prediction Challenge 2019 achieved only low out-of-sample R^2^ when predicting fluid intelligence from T1-weighted MRI data^83^. This discrepancy is likely attributable to the fact that, unlike in the present study, the fluid intelligence scores provided by the competition organizers had been pre-residualized on several covariates—including brain volume, highest parental education, and parental income—all of which are known to be associated with child cognitive ability^12,84^. Residualizing these measures likely removed genuine variance in fluid intelligence from the original scores. Consequently, the proportion of variance that could be explained by brain measures was substantially reduced, helping to clarify why prediction performance in that competition was limited.

### Limitations and future directions

Although our study revealed comprehensive associations between adolescent brain structure and cognitive ability, a key limitation is that we did not incorporate functional brain data. Consequently, we cannot draw conclusions regarding brain activity, an important question we plan to address in future research. Furthermore, our findings are based solely on the Destrieux atlas. While we selected this atlas for its high anatomical resolution and sulco-gyral based delineation, future studies should test the robustness of our findings by using additional parcellation schemes, such as the 68-region Desikan-Killiany (DK) atlas^85^. Additionally, we used DTI and RSI measures provided by the ABCD group to assess gray matter microstructure. These measures were not derived from DWI models specifically optimized for gray matter, such as NODDI^17^, which may provide more accurate quantification despite being computationally more complex.

In this study, we investigated the association between brain structure and cognitive ability using a large, representative adolescent sample and robust large-scale computational models. Our findings provide a foundation, methodology, and guidance for future developmental brain-behavior association studies. Building on this work, we intend to employ brain structural measure selection and develop advanced computational models to characterize adolescent cognitive development in greater depth. These models could aid in identifying atypical cognitive development and offer crucial methodological support for other studies exploring the neural underpinnings of cognitive development.

## Methods

### Participants

The full ABCD 5.1 release cohort includes 27,595 scans of 11,868 participants across 22 sites^39^. The subject selection and exclusion process is shown in **Fig. 7A**. First, we selected brain images that passed quality control^40^ and excluded participants with neurological or psychiatric conditions, as well as observations with missing cognitive scores or brain structural measures. Following this, participants with outliers in cognitive scores or whole-brain structural measures (> 3 standard deviations (*SD*) above or below the mean) were excluded^86^. As depicted in **Fig. 7A**, this process yielded a final cohort of 8,543 participants (6,742 females; 13,992 total scans) aged 9 to 15 years. Because the ABCD dataset is longitudinal, some participants had multiple scans. To remove confounds from repeated scans from the same subject, we randomly selected one scan per participant in each bootstrap iteration for our cross-sectional analyses.

**Fig. 7.**
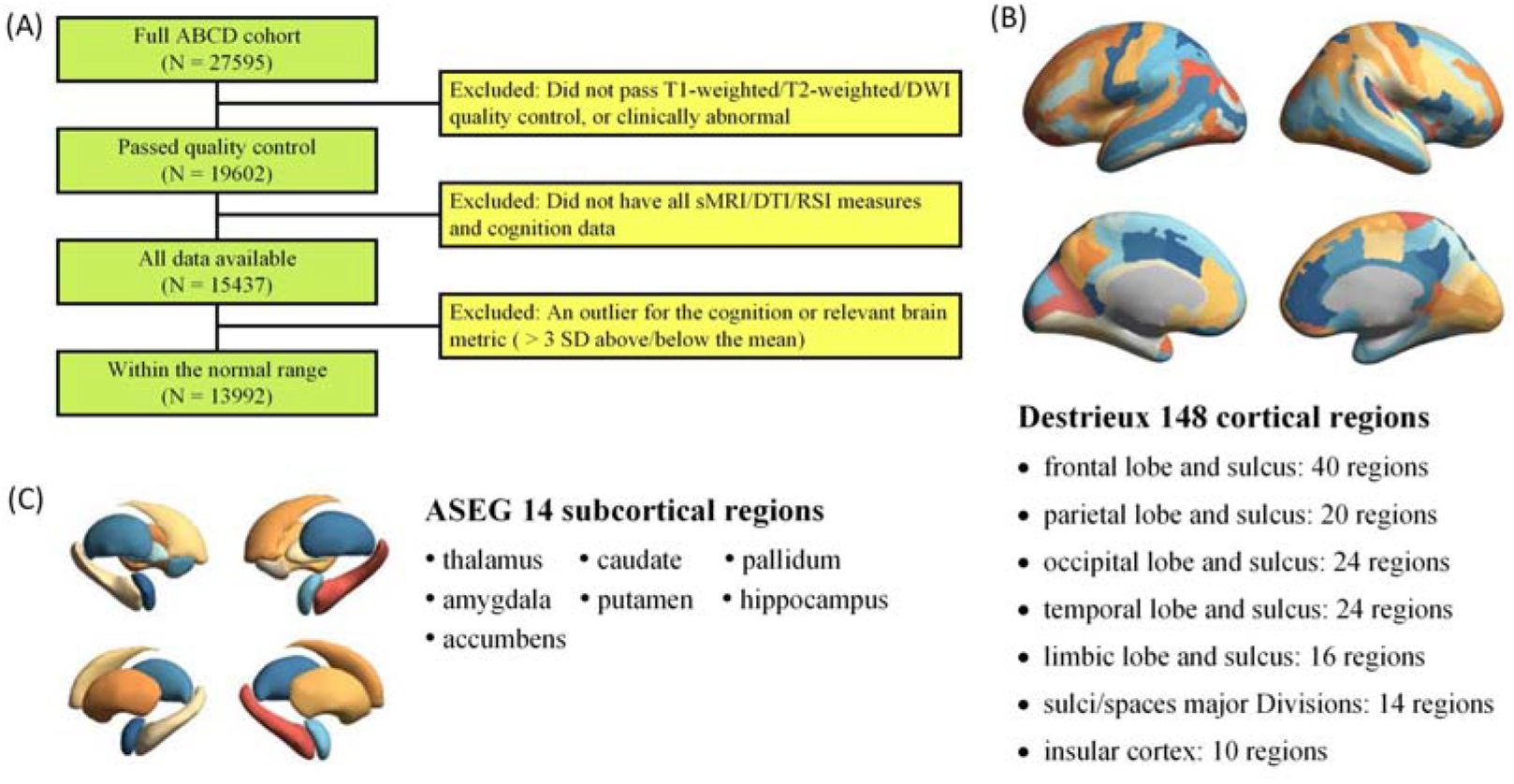
Overview of data preprocessing. (A) Subject selection and exclusion process. (B) 148 cortical regions based on the Destrieux atlas. (C) 14 subcortical regions based on the Automated Subcortical Segmentation (ASEG) atlas.

### Imaging acquisition and processing

We employed three types of brain structural imaging data, comprising T1-weighted, T2-weighted, and DWI recordings, preprocessed by the ABCD group^40^. Structural MRI data (T1w and T2w) were preprocessed by gradient warp correction, bias field correction, and resampling to 1 mm isotropic resolution. DWI data were preprocessed by eddy current distortion, motion correction, B0 distortion correction, gradient warp correction, and resampling to 1.7 mm isotropic resolution. We derived 16 region-wise brain structural measures from these modalities using FreeSurfer 7.1.1 and FSL 5.0.2.2^40^. These measures included 6 sMRI measures: cortical surface area (CA), cortical thickness (CT), sulcal depth (SD), gray matter volume (VOL), T1w density (T1w), and T2w density (T2w). Additionally, there were 4 DTI-based measures: fractional anisotropy (FA), mean diffusivity (MD), longitudinal diffusivity (LD), and transverse diffusivity (TD). Decreased FA and increased LD, MD, and TD were considered to indicate disruptions in white matter structure, such as reduced fiber integrity, smaller axon size, and lower myelination^87,88^. Further, there were 6 RSI measures: hindered normalized directional (HND), hindered normalized isotropic (HNI), hindered normalized total (HNT), restricted normalized directional (RND), restricted normalized isotropic (RNI), and restricted normalized total (RNT). Notably, RSI, an advanced technique, offers enhanced accuracy in depicting tissue microstructure compared to DTI^67,68^. Three measures were calculated for cortical regions only: cortical surface area, cortical thickness, and sulcal depth. In the preprocessing stage, brain cortex parcellation (**Fig. 7B**) was performed using the Destrieux atlas with 148 regions of interest (ROIs), and delineation of subcortical regions (**Fig. 7C**) was achieved using the ASEG atlas with 14 ROIs^89,90^.

### Whole-brain structural measures

We constructed whole-brain structural measures using three complementary approaches: regional structure, structural connections, and graph theoretical hub features (**Fig. 1**). In the following sections, “region” refers to the regional brain structure, “connection” refers to the strength of a structural connection based on individual MSNs, and “hub” refers to the graph theoretical properties of ROIs based on MSNs. As previously mentioned, each of the 148 cortical regions had 16 regional brain structural measures, while each of the 14 subcortical regions had 13. This resulted in a total of 2,550 (*i*.*e*., 148 × 16 + 14 × 13 = 2,550) region-wise inputs, encompassing sMRI, DTI, and RSI features.

For connection-wise inputs, we used MSNs to quantify structural connections between each pair of brain regions, which have been linked to similarities in cytoarchitecture, gene expression, and fiber connection strength^21,91^. Specifically, after normalizing the individual morphometry measures across ROIs, we calculated the connections between cortical regions using Pearson correlation based on the 16 regional structural measures. For connections involving subcortical regions, we used 12 of the 13 available measures; the subcortical volume measure was excluded due to inconsistencies between cortical and subcortical volume calculations. Calculating these individual MSNs for all 162 ROIs (*i*.*e*., 148 cortical + 14 subcortical) yielded 13,041 (*i*.*e*., 162 × (162 - 1) / 2 = 13,041) connection-wise inputs. Further details can be found in **Supplementary Materials S2**.

To characterize each brain hub within individual MSNs, we calculated the six most established graph theoretical hub measures^41^. These measures included strength (degree), clustering coefficient, local efficiency, betweenness centrality, within-module degree, and participation coefficient. Following the approach of Seidlitz et al. (2018), all graph theoretical measures were calculated at multiple connection densities (from 10% to 40% in 5% increments). For each density, weighted graphs were created by setting connection values below the threshold to 0. Results were highly similar across all connection densities, with correlations ranging from 0.41 to 0.55. Because the 35% connection density showed the best performance in a basic correlation analysis with cognitive scores, we focus on the results for this density in the main text; results for all other densities are provided in **Supplementary Materials S3, Supplementary Fig. 6**, and **Supplementary Table 3**. This process yielded 972 (*i*.*e*., 162 × 6 = 972) MSN-based inputs capturing hub-wise characteristics. In total, we extracted a comprehensive set of 16,563 (*i*.*e*., 2,550 + 13,041 + 972 = 16,563) structural inputs encompassing region, connection, and hub characteristics.

### Cognitive measures

To assess cognitive ability, we used seven subtest scores from the ABCD dataset, excluding those with missing data^92^. Some subtests were not collected at certain longitudinal time points due to various factors, including the COVID-19 pandemic. The remaining subtests were vocabulary (PV), attention (Flanker), reading (ORR), processing speed (PCPS), episodic memory (PSM), visuospatial accuracy (LMT Acc), and visuospatial efficiency (LMT EFF). These subtests measure crystallized cognitive ability (vocabulary, reading), fluid cognitive ability (attention, processing speed, episodic memory), and visuospatial ability (visuospatial accuracy and efficiency)^93,94^. We performed a Principal Component Analysis (PCA) on these seven subtests and used the first extracted component as an estimate of general intelligence (*g*), a common practice in the field^26,49^. In total, our cognitive assessment included seven ability-specific scores and the composite *g* score. Although the full ABCD battery includes 12 cognitive subtests, the *g*-factor derived from our 7 subtests was highly correlated with a *g*-factor derived from the complete set of 12 (*r* = 0.89). Further details are provided in **Supplementary Materials S4**.

### Large-scale computational model for investigating associations

We used a large-scale computational model combined with 1,000 family-specific bootstraps (**Fig. 1**) to investigate the association between 16,563 brain structural inputs and our eight measures of cognitive ability. This model was adapted from the method developed by Spreng et al. (2020), with several modifications for the present study. Specifically, the original method calculated independent Pearson correlations between microstructural and behavioral measures over 100 bootstrap iterations, selecting significant associations based on 5–95% confidence intervals (CIs). However, we posited that brain structural measures are not independent but rather complement each other and collectively contribute to cognitive ability. Therefore, we employed a large-scale multivariate regression model instead of independent Pearson correlations, as shown in **Equation 1**, allowing the brain structural measures to control for one another and provide a more integrated analysis.

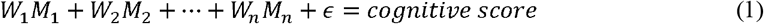

where *M*_*i*_ is the *i*th brain structural input *W*_*i*_ is the corresponding parameter, and ϵ is the error term. We improved the model’s robustness and removed subject and family confounds by introducing a random optimization mechanism and 1,000 family-specific bootstraps, as described later in this section. The model was also adjusted for sex, age, site, and a proxy measure of total brain volume (PBV), consistent with Karama et al. (2011). Here, PBV refers to the total brain volume excluding the cortical and subcortical structures used in our analysis. We used PBV instead of total brain volume because our structural measures already contain cortical and subcortical information; controlling for a total brain volume metric that includes these measures would implicitly correct for our effects of interest.

Given that there were three types of structural brain inputs (region, connection, and hub) and eight distinct cognitive measures, we constructed 24 separate models to investigate the associations. Taking the model for region-wise brain structure and general intelligence (*g*) as an example, we fed 2,550 regional structural inputs into the model to estimate *g*. Following 1,000 bootstraps, we generated 1,000 parameters (*W*_*i*_) for each regional input (*M*_*i*_) to assess its significance in associating with *g*. An input was considered significant only if it passed Bonferroni correction (mean parameter value tested against zero across all regional inputs) and exhibited statistical significance based on 5–95% bootstrap confidence intervals (meaning all parameter values were either non-positive or non-negative)^45,95,96^. The association value for a significant input was then defined as the absolute value of the mean of its 1,000 parameters; the association values for non-significant inputs were set to 0. Using this approach, we obtained association values for all 16,563 structural inputs across all eight cognitive measures. As illustrated in **Fig. 1**, we investigated both specific and summary associations. A specific association is the individual value for a single structural input. A summary association was calculated for region-and hub-wise inputs in two ways: (1) by summing the association values for a given feature (like cortical thickness) across all ROIs, or (2) by summing the association values for a given ROI across all of its features. Summary associations were not calculated for connection-wise results because each connection was represented by a single value, and aggregating these pairwise associations across ROIs would not be meaningful for our analysis.

Our model incorporates several techniques to ensure robustness and validity. First, we used a LASSO method to select relevant features from the large pool of inputs and to mitigate issues arising from the *N/P ratio* problem common in machine learning^81,82^. Second, we employed a feature random optimization strategy, which introduced variability in the order of parameter computation across 1,000 bootstraps to reduce collinearity effects and increase model robustness. Third, we used a family-specific bootstrap to eliminate family-related confounds by ensuring that no two participants from the same family were included in any single bootstrap. Fourth, the large number of bootstraps (1,000) provided a stable basis for our strict feature selection process. Finally, we calculated the coefficient of determination (*R*^*2*^) between the estimated and observed cognitive scores. Although not an out-of-sample validation, these *R*^*2*^ results serve to confirm the effectiveness of our model.

### Large-scale computational model for investigating temporal stability of associations

To investigate the temporal stability of brain-cognition associations during adolescence, we extended the previously described computational model. Based on the model in **Equation 1**, we introduced interaction terms between the brain structural inputs and age to characterize age-related effects on these associations, as shown in **Equation 2**.

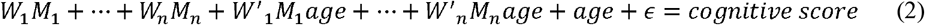

where the absolute value of the parameter *W*’_*i*_ represents the effect of age on the association between brain input *M*_*i*_ and cognitive score. A higher parameter |*W*’_*i*_| means higher instability in the corresponding association during adolescence. All model fitting and significance testing procedures were identical to those used for **Equation 1**. We did not, however, additionally adjust the data for age, as it was already explicitly included as a main effect and in the interaction terms. As before, 24 distinct models were constructed to accommodate the three types of brain structural inputs (region, connection, and hub) and the eight cognitive measures.

## Data availability

The data used in this study were obtained from the ABCD Study, release 5.1, under the terms of a data use agreement. Raw data cannot be shared directly by the authors but are accessible through application to the NIMH Data Archive (NDA). Researchers with approved access can use the provided scripts to reproduce the analyses.

## Code availability

All analysis code is accessible online: https://github.com/JDYan/Brain-Cognition-Association.

## Acknowledgements

Sherif Karama is supported by the Canadian Institutes of Health Research. Alan Evans is supported by CFREF/HBHL. Kirsten Hilger is supported by the German Research Foundation (HI 2185/1-3). Jiadong Yan is supported by the China Scholarship Council. Data used in this study were obtained from the ABCD Study (https://abcdstudy.org), held in the NDA. The ABCD Study is a multisite, longitudinal study designed to recruit over 10000 children aged 9-10 and follow them for 10 years. It is supported by the National Institutes of Health (NIH) and other federal partners under award numbers U01DA041048, U01DA050989, U01DA051016, U01DA041022, U01DA051018, U01DA051037, U01DA050987, U01DA041174, U01DA041106, U01DA041117, U01DA041028, U01DA041134, U01DA050988, U01DA051039, U01DA041156, U01DA041025, U01DA041120, U01DA051038, U01DA041148, U01DA041093, U01DA041089, U24DA041123, and U24DA041147. A complete list of participating sites and investigators is available at https://abcdstudy.org/consortium_members/. The data used in this study were from ABCD Release 5.1.

## Author contributions

Jiadong Yan: Methodology, Software, Formal analysis, Investigation, Writing–original draft, Writing–review & editing, Visualization. Yasser Iturria-Medina: Conceptualization, Methodology, Software, Formal analysis, Investigation. Gleb Bezgin: Methodology, Software, Visualization, Validation. Paule Joanne Toussaint: Methodology, Investigation, Writing–original draft. Ke Xie: Methodology, Visualization. Liang He: Methodology, Visualization. Judy Chen: Validation, Writing–original draft. Kirsten Hilger: Methodology, Validation, Writing–original draft. Erhan Genç: Methodology, Validation, Writing–original draft. Alan Evans: Conceptualization, Funding acquisition, Supervision. Sherif Karama: Conceptualization, Methodology, Investigation, Writing–original draft, Writing–review & editing, Funding acquisition, Supervision.

## Competing interests

The authors declare no competing interests.

## Notes

### Competing Interest Statement

The authors have declared no competing interest.

### Summary of Updates

New version with new results and discussion

